# MELO-ED: learning locality-sensitive multi-embeddings for edit distance

**DOI:** 10.1101/2025.11.23.689944

**Authors:** Xin Yuan, Ke Chen, Ajmain Yasar Ahmed, Mingfu Shao

## Abstract

Edit distance is a fundamental metric for quantifying similarity between biological sequences, but its high computational cost limits large-scale applications. Previously, we proposed learned locality-sensitive bucketing (LSB) functions that achieved superior performance and efficiency compared to classical seeding methods for identifying similar and dissimilar sequences. However, each component of an LSB function is represented as a one-dimensional hash value that can only be compared for identity, which constrains the method’s accuracy. Here, we introduce MELO-ED, a multi-embedding locality-sensitive framework that upgrades each hash value to a higher-dimensional embedding capable of efficiently approximating edit distance. MELO-ED employs a Siamese convolutional neural architecture that learns complementary embeddings capturing both global sequence context and fine-grained edit operations. By integrating locality-sensitive bucketing with multi-embedding representations, MELO-ED achieves near-perfect accuracy without increasing the number of buckets required. Leveraging mature indexing methods in the embedding space, MELO-ED transforms time-consuming edit distance computations into scalable similarity searches across massive genomic databases. Comprehensive evaluations on simulated DNA sequences and real barcode datasets demonstrate that MELO-ED outperforms both traditional alignment-free methods and contemporary machine learning approaches, including our previously developed learned LSB functions. These results establish MELO-ED as a state-of-the-art framework for fast and accurate classification of similar and dissimilar sequences. MELO-ED is available at https://github.com/Shao-Group/MELO-ED.

## 1 Introduction

Many fundamental tasks in sequence analysis require identifying all pairs of similar sequences within a large dataset or across multiple datasets. Examples include constructing overlap graphs from long reads, a critical step in genome assembly [2, 20, 11, 39, 23, 15, 22, 21, 13], building phylogeny trees from sequencing data [14, 30, 36, 9, 40], and classifying metagenomic datasets [41]. This problem is computationally challenging for two major reasons. First, edit distance is typically used to define sequence similarity/dissimilarity, as it provides a biologically meaningful model of evolutionary mutations, but it is expensive to compute and is believed not to admit a strongly subquadratic-time algorithm [1]. Second, the naive all-vs-all comparison approach is prohibitively costly for datasets that often contain millions of sequences.

Existing efficient methods for this problem can be broadly classified into two categories. The first category relies on the central idea of *bucketing*, where each sequence is assigned into one or multiple buckets, and all-vs-all comparisons are carried out only within individual buckets. Seeding and sketching methods, such as minimizers [32, 37, 33, 25], syncmers [12], and strobemers [34, 24, 35], follow this paradigm by comparing two sequences only if they share at least one *k*-mer or seed. In general, the goal is to place similar sequences into the same bucket while keeping dissimilar sequences in separate ones. Locality-sensitive hashing (LSH) formalizes this principle. A family of hash functions is said to be (*d*_1_*, d*_2_)-sensitive for a distance function *d*(·, ·) if two sequences *x* and *y* collide (i.e., fall into the same bucket) with high probability when *d*(*x, y*) ≤ *d*_1_, and with low probability when *d*(*x, y*) ≥ *d*_2_. An LSH is ungapped if *d*_1_ = *d*_2_ − 1, and gapped if *d*_1_ *< d*_2_ − 1. Designing LSH functions for edit distance has been a long-standing challenge. A recent breakthrough, Order Min Hash (OMH), was proved to be a gapped LSH [26]. However, achieving sufficiently high collision probability for pairs with edit distance at most *d*_1_ while maintaining low collision probability for pairs with distance at least *d*_2_ requires a large gap between *d*_1_ and *d*_2_, which limits OMH’s practical utility.

On the other hand, a simple argument shows that perfect, deterministic LSH functions for edit distance cannot exist. Suppose there were a function *f* which satisfies *f* (*x*) = *f* (*y*) with probability 1 for every pair of sequences *x* and *y* with edit(*x, y*) ≤ *d*_1_. Then, for any pair *x* and *z* with edit(*x, z*) ≥ *d*_2_, one can construct a series of intermediate sequences *y*_1_*, y*_2_, · · · *, y_t_* such that edit(*x, y*_1_) = edit(*y*_1_*, y*_2_) = · · · = edit(*y_t_, z*) = 1 ≤ *d*_1_, which would imply *f* (*x*) = *f* (*z*) with probability 1 as well. Motivated by this inherent limitation, we introduced *locality-sensitive bucketing* (LSB) functions [5], as a generalization of LSH. An LSB function *f* maps each sequence into multiple buckets, so that if *d*(*x, y*) ≤ *d*_1_, then *f* will map them into at least one shared bucket, i.e., *f* (*x*) ∩ *f* (*y*) ≠ ∅; and if *d*(*x, y*) ≥ *d*_2_, then they are mapped to disjoint sets of buckets, i.e., *f* (*x*) ∩ *f* (*y*) = ∅. Unlike perfect LSH functions, we proved that such LSB functions do exists! We designed LSB functions for several distance parameters (*d*_1_*, d*_2_), including (1, 2), (1, 3), (3, 5), and (2*r* − 1, 2*r* + 1). However, constructing perfect LSB functions for arbitrary *d*_1_ and *d*_2_ with a small number of buckets per sequence remains challenging. To address this, we proposed to learn LSB functions from data, representing the bucketing functions as a neural network with trainable parameters [43]. The learned LSB functions achieve near-perfect accuracy for several (*d*_1_*, d*_2_) settings, matching the corresponding theoretical results, but for others, such as (2, 3) and (5, 6), the accuracy at the boundary distances is considerably lower, leaving substantial room for improvement.

The second category of approaches follows a two-step framework: sequences are first embedded into a latent space, after which LSH is applied within the embedding space. Normed spaces such as the Hamming or Euclidean spaces are commonly used because efficient LSHs are well established for them [16, 8]. Considerable effort has been devoted to embedding sequences under edit distance into such normed spaces with low distortion [31], where distortion is defined as the supremum of the ratio between the embedding distance and the true edit distance, over all sequence pairs. However, existing embeddings suffer from large distortion; for instance, the CGK embedding maps sequences into Hamming space with distortion of *O*(*k*^2^) where *k* denotes the edit distance [3]. Lower-bound results further show that edit distance cannot be embedded into *l*_1_ norm with distortion better than Ω(log *N*), where *N* is the sequence length [19]. A number of heuristic embedding methods have also been proposed, including FFP [38] and Tensor Sketching [18, 17], which generates embeddings by counting or profiling fixed-length substrings or subsequences. Machine learning-based approaches have also been explored, such as GRU [44], CNNED [7], Bio-kNN [4], and NeuroSEED [6], which employ various neural architectures ranging from gated recurrent units to convolutional neural network and transformers to encode sequences into vector representations. Although these methods demonstrate empirical improvements in specific applications, they remain fundamentally constrained by the aforementioned theoretical limitations: all of them map each sequence to a single vector, which inherently limits how well they can approximate edit distance in any normed space.

In this work, we combine the strengths of the above two paradigms by extending traditional single-vector embeddings into multi-embeddings. Formally, a multi-embedding function *g* maps a sequence *s* to a tuple of *k* vectors, (*g*_1_(*s*)*, g*_2_(*s*), · · · *, g_k_*(*s*)) in a latent space. The goal is to ensure the following locality-sensitive behavior: if two sequences *s* and *t* satisfy edit(*s, t*) ≤ *d*_1_, then they are close in at least one embedding dimension, i.e., there exists an index *i* such that l*g_i_*(*s*) − *g_i_*(*t*)l *< δ* for a predefined distance threshold *δ* in the latent space; and conversely, if edit(*s, t*) ≥ *d*_2_, then they are far apart in all embedding components, i.e., l*g_i_*(*s*) − *g_i_*(*t*)l *> δ* for all 1 ≤ *i* ≤ *k*. Intuitively, multi-embedding mitigates the limited expressiveness of hash functions in LSH or LSB, where two sequences either collide or do not, while also circumventing the theoretical constraints of single-vector embeddings by distributing the representational burden of a faithful embedding across multiple functions—although individual embedding components may still exhibit large distortion, strong overall performance can be achieved as long as they complement one another.

We train such multi-embedding functions for arbitrary *d*_1_ and *d*_2_, a method we call MELO-ED (pronounced melody), stands for <u>M</u>ulti-<u>E</u>mbedding-based <u>Lo</u>cality-sensitive bucketing for <u>E</u>dit <u>D</u>istance. Our results show that MELO-ED achieves substantially higher accuracy than single-embedding functions. Moreover, for (*d*_1_*, d*_2_) settings where the learned LSB functions previously exhibited low accuracy, MELO-ED delivers significant improvement and in several cases attains near-perfect performance. When combined with existing nearest neighbor indexes in the latent space, MELO-ED provides drastically improvements over existing hashing- and embedding-based techniques for large-scale sequence comparison, as well as in an application that involves barcode classification.

## 2 Multi-Embedding: Definition, Learning, and Comparison

### 2.1 Problem Formulation

Let *g* be a function that maps a string *s* of length *N* over alphabet Σ = {*A, C, G, T* } to *k* vectors, (*g*_1_(*s*)*, g*_2_(*s*), · · · *, g_k_*(*s*)), in the Euclidean space, i.e., *g_i_*(*s*) ∈ R*^m^*, 1 ≤ *i* ≤ *k*. We define such a function *g* to be a (*d*_1_*, d*_2_)-locality-sensitive multi-embedding (LSME) function, *d*_1_ *< d*_2_, if for every two strings *s* and *t* over Σ, if edit(*s, t*) ≤ *d*_1_ then min_1≤_*_i_*_≤_*_m_* l*g_i_*(*s*) − *g_i_*(*t*)l_2_ ≤ *δ*, and if edit(*s, t*) ≥ *d*_2_ then min_1≤_*_i_*_≤_*_m_* l*g_i_*(*s*) − *g_i_*(*t*)l_2_ *> δ*, where edit(*s, t*) denotes the edit distance between *s* and *t* and l · l_2_ denotes the Euclidean distance. Intuitively, an LSME function guarantees that at least one pair of embedding vectors will be close for similar strings and all pairs of embedding vectors will be distant for dissimilar strings. See Fig. 1 for an illustration. A (*d*_1_*, d*_2_)-LSME function is parameterized by *N*, *k*, *m*, and *δ*.

**Figure 1:**
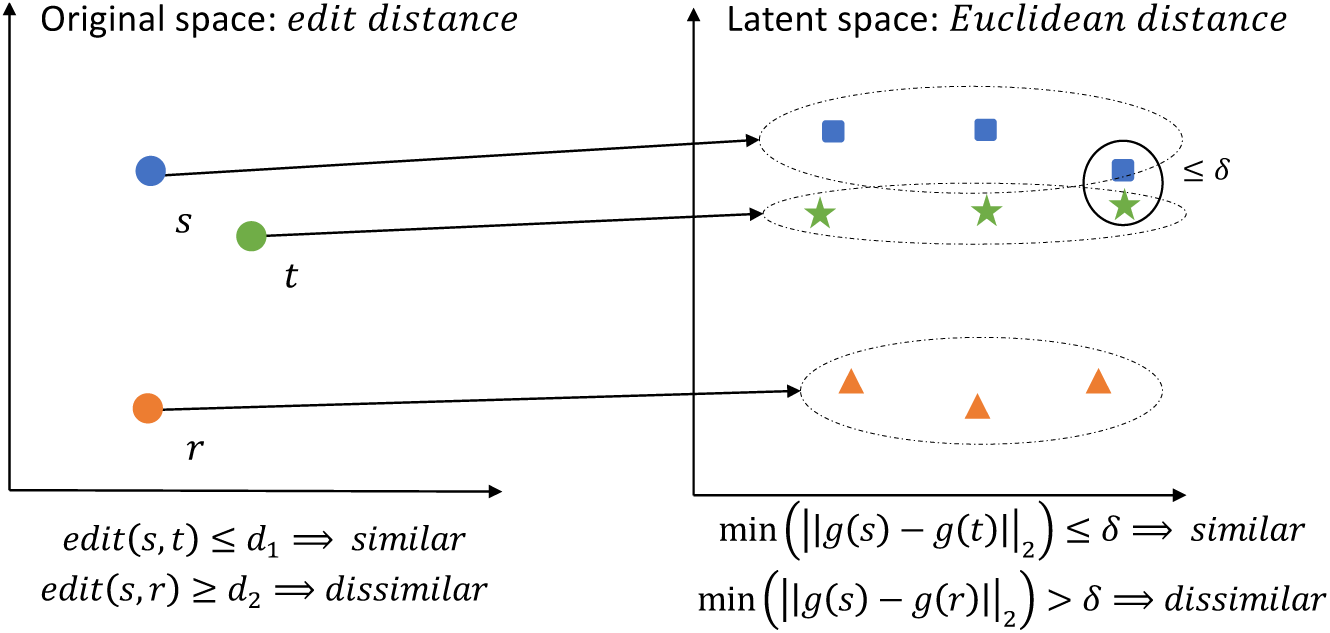
Left: original space measured by edit distance. Right: latent space measured by Euclidean distance.

### 2.2 LSME Representation: Inception CNN

We represent an LSME function *g* as a neural network with learnable parameters. The architecture of the neural network uses Inception modules to capture multiscale patterns within DNA sequences, allowing the model to learn both short- and long-range dependencies.

Fig. 2 illustrates the structure of a two-layer inception model. The input sequence *s*, of length *N*, is one-hot encoded and represented as a matrix *s* of shape [1, 4*, N*], where the original channel is 1 and the middle 4 dimensions correspond to {*A, G, C, T*} separately. Each Inception CNN layer consists of *l* CNN units (*l* = 6), which apply parallel convolutional filters with different kernel sizes to extract a diverse set of features from the sequence. The kernel size is defined as *c* × (*l* + 1) × 4, where *c* denotes the channel numbers and (*l* + 1) × 4 denotes the window size of each kernel. Each CNN unit in the Inception layer is activated by a rectified linear unit, *ReLU* (·), followed by a max-pooling layer that selects the maximum value within a window of size *p*. The padding shape for each CNN unit’s output is set to ensure that the merged output is feasible. The output of the Inception CNN layer is composed across all channels, resulting in a matrix of dimension [*l* × *c,* 4*, N* − *p*]. Two or more such Inception CNN layers can be stacked as needed. At the bottom of the model is a linear layer with the input from the final Inception CNN layer, which is flattened by a Flatten layer. The linear layer maps the flattened feature representations to a matrix of size *k* × *m*, giving the desired *k* embedding vectors of dimension *m*.

**Figure 2:**
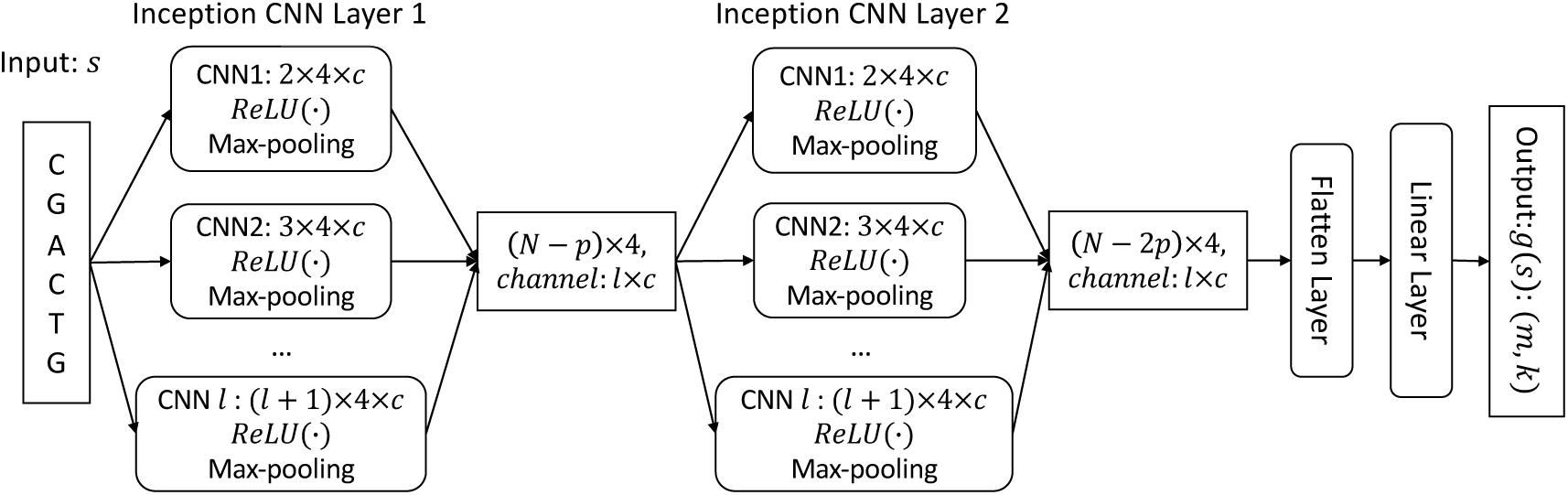
Structure of a two-layer inception convolutional neural network that represents an LSME function.

### 2.3 Training Framework: Loss Function and Siamese Network

We develop a framework that can train an (*d*_1_*, d*_2_)-LSME function with chosen *N*, *k*, *m*, and *δ*. Recall that an LSME function *g* is defined over two sequences. Hence, the training data consists of pairs of sequences {(*s, t*)}. The label *y* of a training sample (*s, t*) is determined based on *d*_1_ and *d*_2_: we define *y* = 1 if *edit*(*s, t*) ≤ *d*_1_ and *y* = −1 if *edit*(*s, t*) ≥ *d*_2_. On one training sample the contrastive loss is defined as:

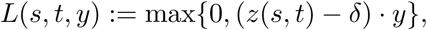

Where

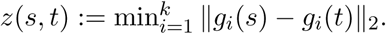

The above contrastive loss enforces the desired locality-sensitive properties. To see this, consider a similar pair (*s, t*) with *y* = 1, i.e., *edit*(*s, t*) ≤ *d*_1_; it is desirable to have *z*(*s, t*) ≤ *δ*. One can verify that if *z*(*s, t*) *> δ* then the loss *L*(*s, t, y*) is strictly positive and if *z*(*s, t*) ≤ *δ* the loss is 0. Similarly, for a dissimilar pair (*s, t*) with *y* = −1, we have that the loss *L*(*s, t, y*) is 0 if *z*(*s, t*) ≥ *δ* and strictly positive if *z*(*s, t*) *< δ*.

To enable training with pairs, we use the Siamese neural network. See Fig. 3. The Siamese neural network contains two copies of the LSME function *g*, but they share model Parameters and remain identical throughout the training. The function *g* maps *s* and *t* into *k* embedding vectors separately, and the loss is calculated as described above.

**Figure 3:**
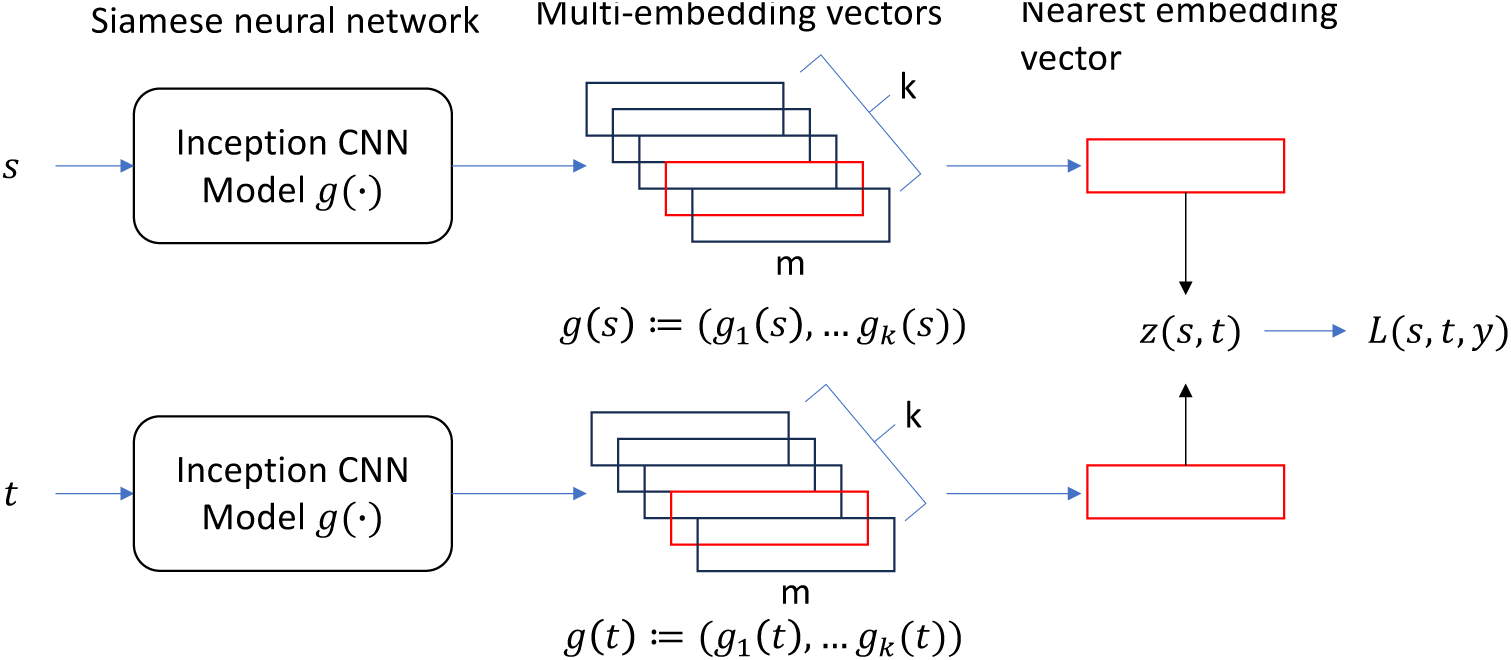
Siamese neural network trained by a hinge contrastive loss function *L*(*s, t, y*).

### 2.4 Training Data and Evaluation Metric

The LSME functions are trained solely on simulated data. The training set includes pairs (*s, t*), generated randomly as follows: starting with a random sequence *s* of length *N*, a series of edits is applied to produce *t*. Each edit (substitution, insertion, or deletion) occurs with a one-third probability at a randomly chosen position. Afterwards, *t* is either padded with random characters from the alphabet or truncated to match the length *N* of *s*. The edit distance *d* = edit(*s, t*) is then calculated, and the pair (*s, t*) is grouped according to the value of *d*. The training of the (*d*_1_*, d*_2_)-LSME function uses an active learning approach that only utilizes boundary training data, collecting pairs with an edit distance in the range of *d* ∈ {*d*_1_ − 1*, d*_1_*, d*_2_*, d*_2_ + 1}. This method is motivated by two observations: models at boundaries usually have lower accuracy, and training on boundary data empirically improves accuracy across other edit distances (see Results). Although our training framework can support training of arbitrary LSME functions, in this work, we demonstrate performance and applications for three sequence lengths. For *N* = 16 and *N* = 20, we randomly collect 2,500,000 samples per boundary edit distance. For *N* = 100, we select 500,000 samples from each edit distance, and implement 10-fold cross-validation for each sequence length. All models are trained on an A100 GPU. Training for *N* = 16 and *N* = 20 takes around 9 hours, while training for *N* = 100 takes about 14 hours.

The LSME function is being tested across a wider range of edit distances: for *N* = 20, we examine *d* ∈ {1, 2, · · ·, 15}, and for *N* = 100, *d* ∈ {1, 2, · · ·, 35}. We randomly select 20,000 pairs for each edit distance within the test range. Let (*s, t*) be a test pair with *d* = *edit*(*s, t*), and let *g* be a trained (*d*_1_*, d*_2_)-LSME function (referred to as MELO-ED in the figures). We denote the function *z*(*s, t*) as *z*(*s, t*) := min*^k^* l*g_i_*(*s*) − *g_i_*(*t*)l_2_., which represents the minimum Euclidean distance between the representations *g_i_*(*s*) and *g_i_*(*t*) across the *k* embedding vectors. If *d* ≤ *d*_1_ and *z*(*s, t*) ≤ *δ*, or *d* ≥ *d*_2_ and *z*(*s, t*) *> δ*, then this sample is correctly classified. The accuracy on edit distance *d* is estimated as the fraction of correctly classified samples using the testing data for edit distance *d*. The overall accuracy of the model is estimated as the fraction of correctly classified samples using all testing data across the full range of *d*.

### 2.5 Discussion of Parameters

A (*d*_1_*, d*_2_)-LSME function for sequence length *N* is equipped with 3 key parameters: the number of embedding vectors *k*, the dimension of each embedding feature *m*, and the margin parameter *δ*. While the effect of *k* is analyzed in Section 2.6, here we study how *m* and *δ* influence overall accuracy. Figure 4 shows that increasing *m* consistently enhances performance across various (*d*_1_*, d*_2_) configurations, especially for more challenging decision boundaries, confirming that higher embedding capacity allows for better edit-distance representation; therefore, we choose *m* = 40 for discussing *δ* and the following downstream experiments. Likewise, Figure 5 demonstrates that larger *δ* values produce higher accuracy by enforcing stronger separation between similar and dissimilar sequence pairs in the latent metric space; thus, we adopt *δ* = 10 for discussing *m* and the following downstream experiments.

**Figure 4:**
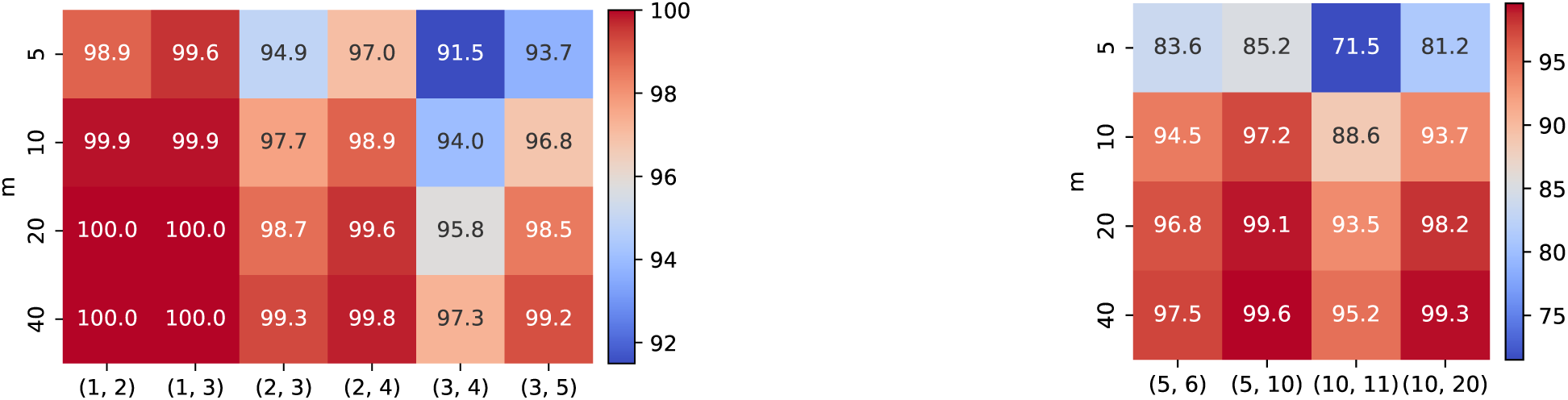
Overall accuracy of MELO-ED on various *m*. Left: *N* = 20, *k* = 20, *δ* = 10. Right: *N* = 100, *k* = 50, *δ* = 10.

**Figure 5:**
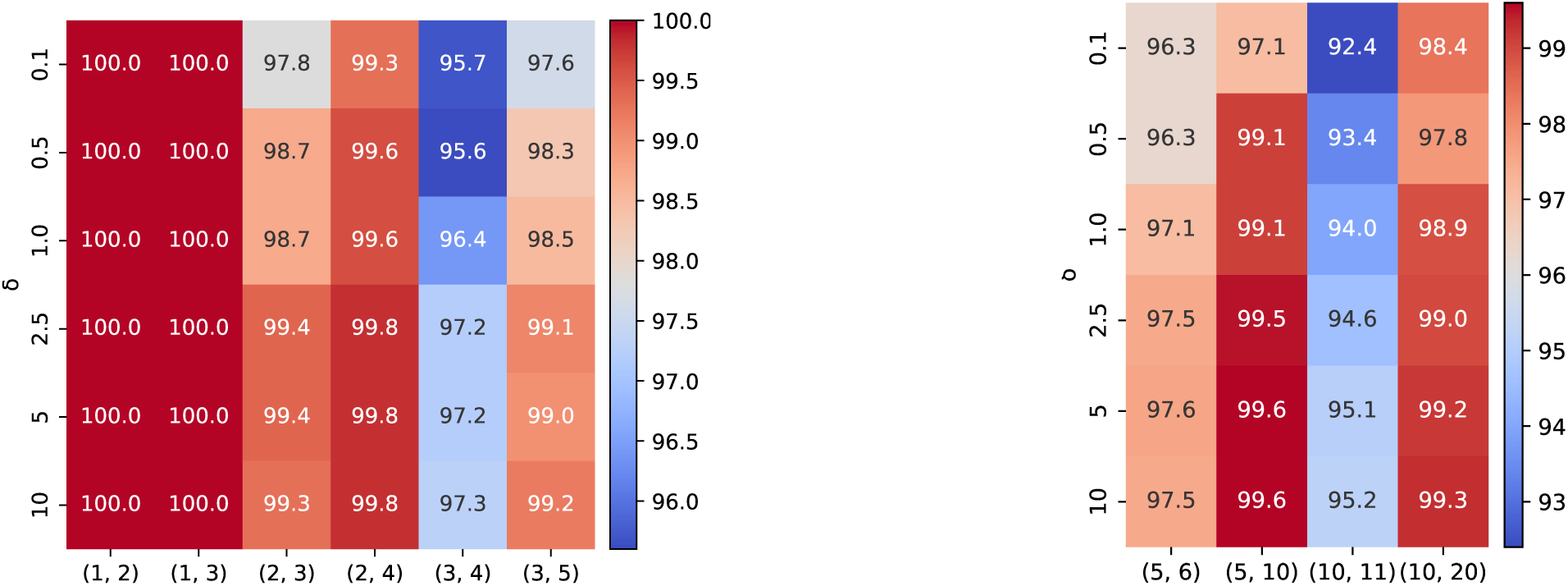
Overall accuracy of MELO-ED on various *δ*. Left: *N* = 20, *k* = 20, *m* = 40. Right: *N* = 100, *k* = 50, *m* = 40.

### 2.6 Comparison with Learned LSB Functions

We evaluate the accuracy of MELO-ED against learned (LSB) functions [43]. Both approaches map a string into *k* vectors in a latent space; however, the LSB function determines similarity based on whether the embedding vectors are identical: two sequences are deemed similar if they share at least one identical vector. In contrast, a LSME function measures similarity using Euclidean distance: two sequences are considered similar if a pair of vectors is within a range smaller than the threshold *δ*. Fig. 6 compares the overall accuracy with varying *k*. MELO-ED consistently outperforms learned LSB functions; the improvements are pronounced for challenging functions such as large *N* or large *d*_1_ and *d*_2_. Note also that MELO-ED achieved very high accuracy across all configurations and nearly perfect accuracy on several cases. Fig. 7 compares the categorized accuracy for individual edit distance with vary *k*. MELO-ED drastically outperforms learned LSB functions on boundary categories, where the latter performs poorly. These results firmly prove the advantages of learning an LSME function.

**Figure 6:**
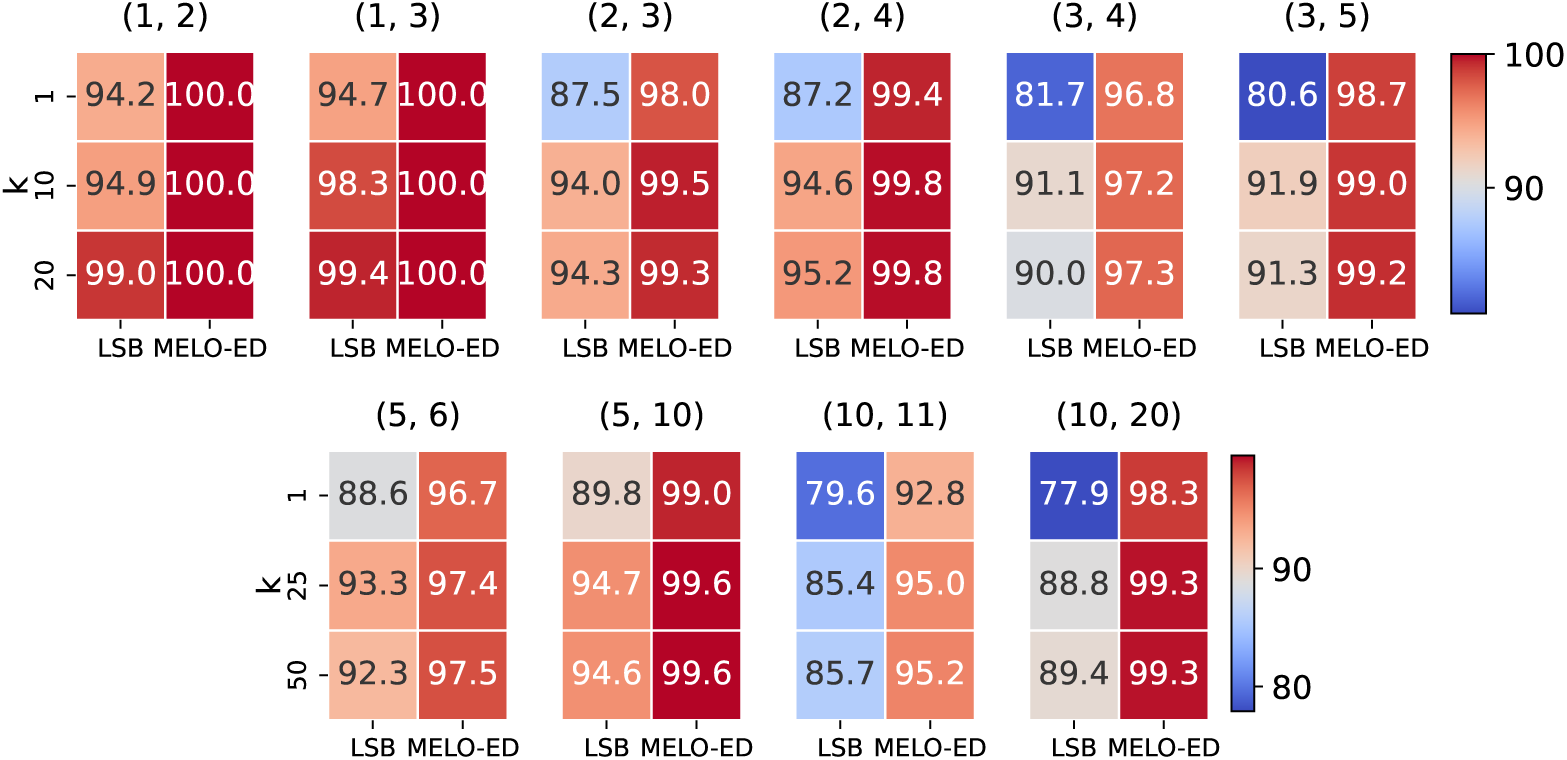
Comparison of learned LSB and MELO-ED under different *k*. Top: *N* = 20, *m* = 40, *δ* = 10. Bottom: *N* = 100, *m* = 40, *δ* = 10.

**Figure 7:**
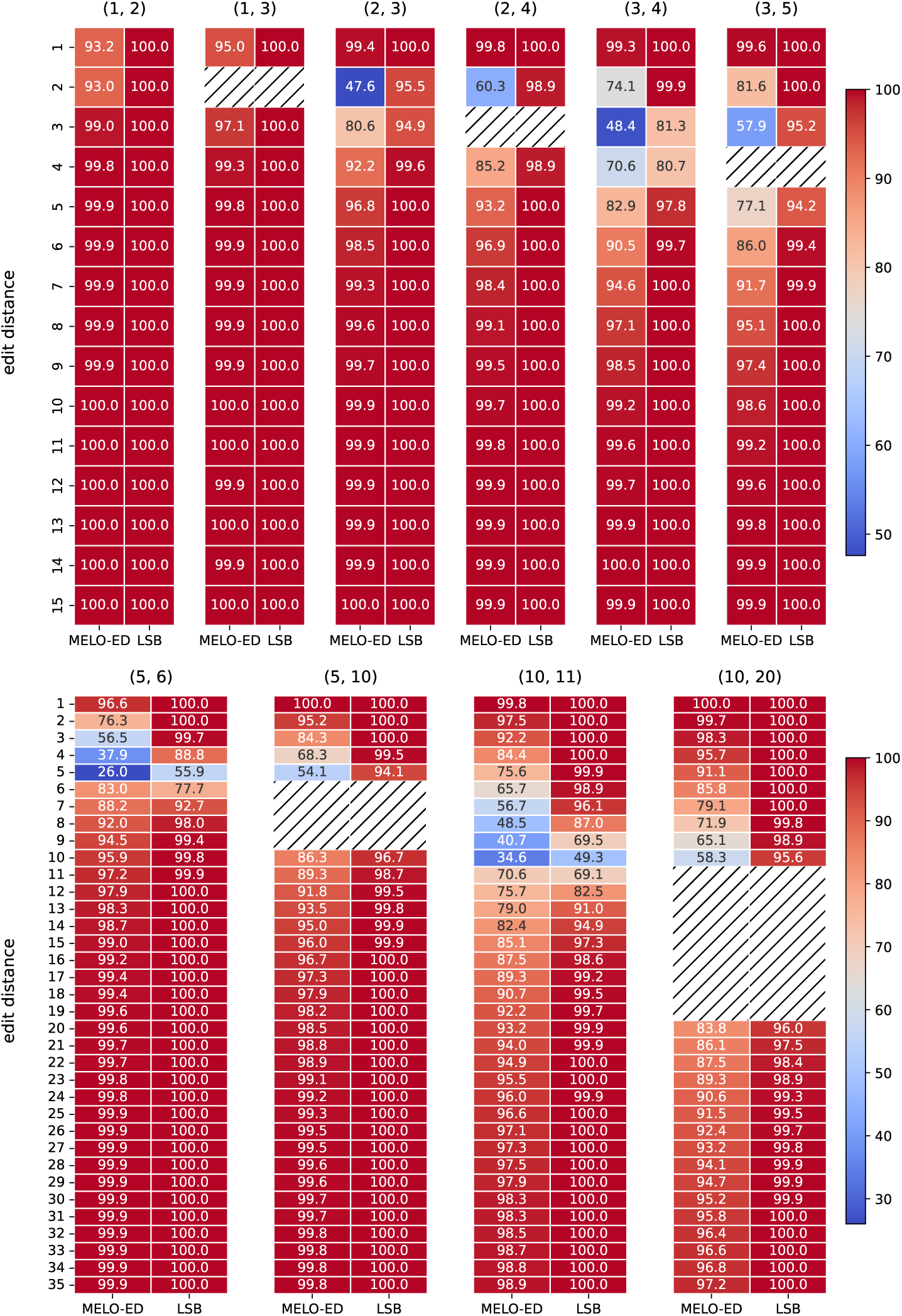
Categorized accuracy of various lengths of sequences. Top: *N* = 20, *m* = 40, *k* = 20, *δ* = 10. Bottom: *N* = 100, *m* = 40, *k* = 50, *δ* = 10.

## 3 Large-Scale Sequence Comparison Using LSME Functions

### 3.1 Indexing Embedding Vectors

A typical application involves large-scale sequence comparison requires to find all sequences in a base set *B* that are within edit distance *d*_1_ from each sequence in a query set *Q* (it could be *B* = *Q* in which case we want to report all pairs of sequences in the set that have edit distances at most *d*_1_). Although the Euclidean distance between two embedding vectors can be calculated more easily than the edit distance between the original sequences, it remains inefficient (or even infeasible) to compute all-vs-all embedding distances for large base and query sets. In order to use our learned embeddings in such applications, we utilize an index for neighbor search, which allows for finding close vectors in the Euclidean space much more efficiently.

There are two types of index available: *K*-nearest neighbor (KNN) search and radius search. The latter is a more natural choice for our case: setting the search radius to the margin *δ* yields all sequences whose embeddings are within *δ* from the query, which, by definition of our model, are sequences within edit distance *d*_1_. The KNN index has the advantage of being more widely-available, and potentially more efficient. It requires an estimation *K* of the number of similar sequences in the base for each query: an under-estimation can miss true positives while an over-estimation requires more verification time to filter out the reported neighbors that are not within the margin *δ*. Since many applications come with a natural choice for *K* (e.g., find the most similar barcode in the whitelist, see Section 4), we also test with a KNN index. In our experiments, for KNN search, we use the Hierarchical Navigable Small World (HNSW) index implemented in Faiss [10]; for radius search, we use the FLANN index [27, 28, 29].

### 3.2 Applying Neighbor Search Index on Testing Data

We apply MELO-ED combined with indexing for neighbor search and compare their accuracy with competing methods. We pick 20,000 pairs of sequences of length *N* = 20 for each edit distance from 1 to 15 from the testing dataset used in Section 2. We collect the first sequences from these 15 × 20, 000 = 3 × 10^5^ pairs to form the query set *Q*, and the second sequences to form the base set *B*. Pairwise edit distances between sequences in *Q* and *B* are computed to build the ground-truth. In a (*d*_1_*, d*_2_)-sensitive search, all methods aim to identify as many pairs with edit distance at most *d*_1_ as possible (i.e., achieve high sensitivity) while keeping the number of reported pairs with edit distance at least *d*_2_ as small as possible (i.e., maintain low false positives or high precision).

For MELO-ED, we build indexes using both HNSW [10] and FLANN [29]. The HNSW index supports only *K*-nearest-neighbor (KNN) search. In our constructed dataset, we set *K* = 1. Note that for many query sequences, even their nearest neighbor in the base set may have an edit distance exceeding the query threshold *d*_1_, which implies that their smallest embedding distance is likely to be greater than the MELO-ED margin *δ*. Therefore, we further filter the returned neighbors by retaining only those whose embedding distances from the query fall within the threshold *δ*, as required by the MELO-ED setup. We emphasize that this filtering step differs fundamentally from filtering by edit distance: first, computing embedding distances is significantly more efficient; and second, it provides a faithful measure of MELO-ED’s intrinsic accuracy, as we are effectively asking “*Should this pair be reported according to MELO-ED?*” rather than “*Is this pair in the ground truth?*” The FLANN index supports radius search, for which we simply set the query radius to match the threshold *δ* of MELO-ED.

We compare MELO-ED with three other widely used methods, each representing a distinct class of approaches. For seeding-based similarity search, we implement the minimizer strategy: for each length-*N* sequence, we select the smallest *k*-mer (under a random ordering) as its seed, and consider two sequences similar if they share a common seed. For a fair comparison, we extract 20 minimizers for each sequence using 20 different random orders, matching the number of embeddings used in MELO-ED. The parameter *k* controls the trade-off between sensitivity and precision of minimizers. We tested a range of sensible *k* values and report two representative cases: one with high sensitivity and one with high precision. The results for other *k* values are provided in the appendix. For embedding-based approaches, we compare against the celebrated CGK embedding [3]. This method randomly embeds sequences from the edit distance space into the Hamming distance space with bounded distortion; that is, sequences that are close under edit distance are expected to remain close under Hamming distance after embedding. For a fair comparison, we generate 20 random CGK embeddings for each sequence, matching the setup of MELO-ED. As with MELO-ED, an index is required for CGK to avoid exhaustive pairwise embedding distance computation. We employ the canonical index for Hamming distance based on the random bit-sampling LSH, as used in [39]. The number of sampled bits *k* controls the balance between sensitivity and precision of the LSH; two representative results are shown in Fig. 8, with additional results included in the appendix. Finally, we also compare against our learned LSB functions [43]. The LSB functions are trained to map each sequence to 20 buckets, matching the number of embeddings per sequence of MELO-ED.

**Figure 8:**
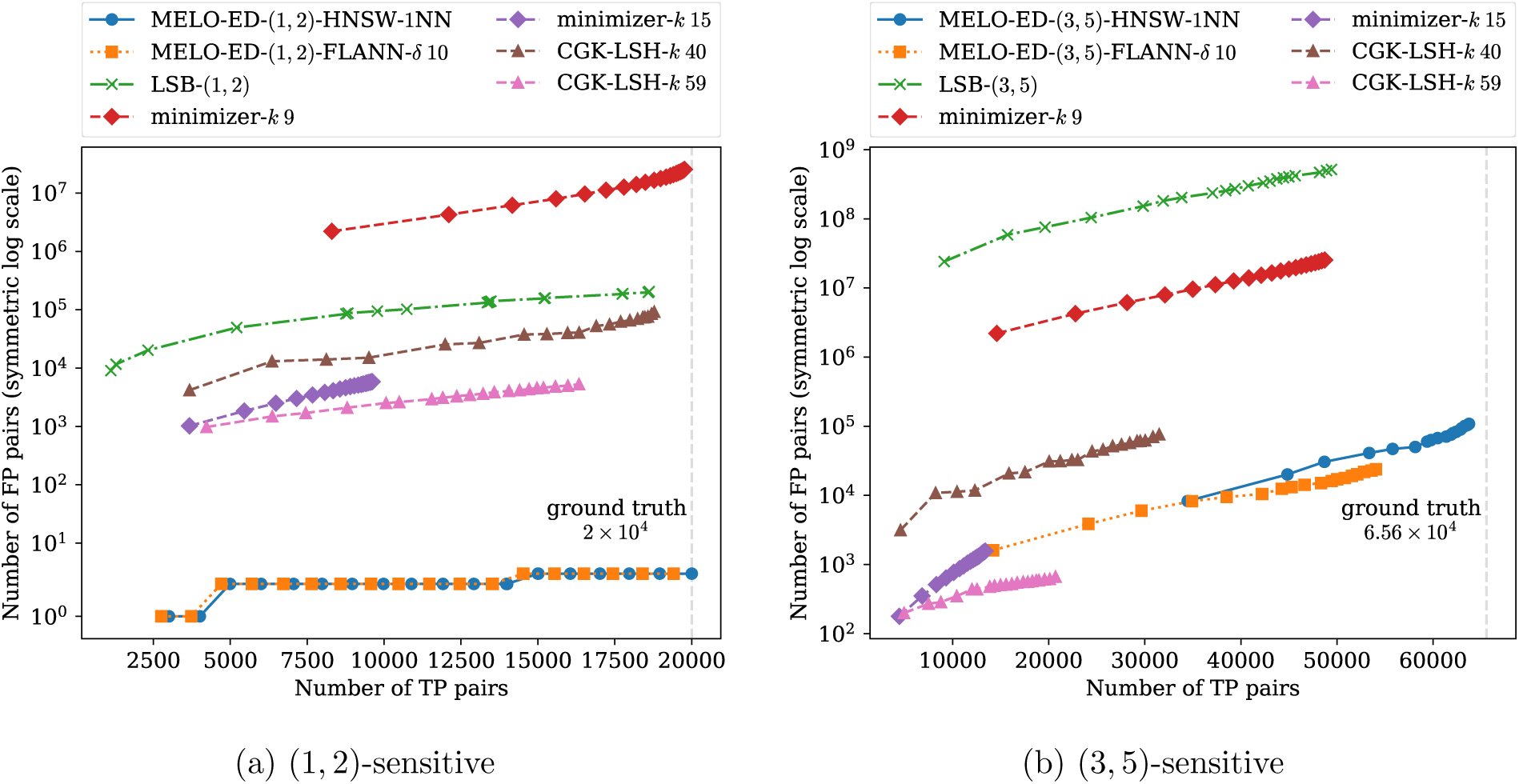
Results of applying a neighbor search index to the embeddings produced by MELO-ED on the test dataset, compared with other methods. A vertical dashed line in each subfigure marks the number of ground-truth similar pairs.

Figure 8 presents the results of different methods evaluated under similarity thresholds (*d*_1_*, d*_2_) = (1, 2) and (3, 5), respectively. Results for other thresholds are included in the appendix. A reported pair is classified as a TP if their ground-truth edit distance is at most *d*_1_; and as an FP if its edit distance is greater than or equal to *d*_2_. Consequently, in a gapped setting (e.g., a (3, 5)-sensitive search), reported pairs with edit distances between *d*_1_ and *d*_2_ are ignored, as they lie in the “don’t care” region defined by the application’s similarity search thresholds. Since each method effectively performs 20 repetitions (20 seeds per sequence for minimizers, 20 buckets for learned LSB, and 20 embeddings for CGK and MELO-ED), we plot each configuration as an interpolated curve with 20 points, where the *i*-th point represents the union of all reported pairs from the first *i* repetitions. As expected, sensitivity increases along with the total number of reported pairs as more repetitions are aggregated.

Across all settings, MELO-ED consistently achieves the highest accuracy, exhibiting nearly perfect recall and a low false-positive rate. In contrast, other methods either suffer from low recall or must tolerate orders-of-magnitude more false positives to approach MELO-ED’s level of sensitivity. The two tested indexes, HNSW and FLANN, produce comparable overall results. However, since both perform *approximate* neighbor searches, it is noteworthy that the false positives they produce differ substantially, suggesting that intersecting results from multiple indexes could potentially further improve precision.

These results further highlight the substantial gains achieved by MELO-ED over LSB models. Although LSB already attains test accuracies exceeding 90% (see Fig. 6), as demonstrated here, even a modest error rate can translate into a large number of incorrect reports in large-scale tasks. The improved accuracy of MELO-ED therefore represents more than an incremental advantage: it provides a meaningful practical benefit by enabling near-perfect recall without requiring costly downstream filtering to eliminate large volumes of false candidates. This advantage is particularly pronounced in gapped similarity settings, where MELO-ED maintains high accuracy even on challenging boundary cases that the LSB models struggle with.

## 4 Application: Barcode Classification

We further evaluate MELO-ED on a real dataset consisting of error-prone cell barcodes. Cell barcodes are synthetic sequences attached to reads to label the cell of origin and are widely used in single-cell sequencing. Similar to the reads themselves, these barcodes may contain errors. When assigning reads to a known list of valid barcodes (the so-called whitelist), erroneous barcodes that cannot be matched are typically discarded, leading to a substantial loss of reads. In our previous work, we showed that learned LSB functions could be used to recover some of these erroneous barcodes (and their associated reads). With the substantially improved accuracy of MELO-ED, we revisit this task to demonstrate its superior classification capability.

We use the dataset from [42], which contains 2,899,770 reads. Among these, 27.3% (790,702 reads) have barcodes that do not match any of the |*W* | = 806 entries in the whitelist. The whitelist *W* serves as the base set, and the 400, 919 unique erroneous barcodes (since multiple reads may share the same erroneous barcode) form the query set *B*. To construct the ground truth, we compute the pairwise edit distances between all barcodes in *W* and *B*. As in Section 3.2, we compare MELO-ED against minimizers, the CGK embedding, and learned LSB functions. For (*d*_1_*, d*_2_) = (1, 2) and (3, 5), the TP–vs-FP curves for all methods, parameterized by the number of repetitions included, are shown in Fig. 9. Additional results for other similarity thresholds are provided in the appendix.

**Figure 9:**
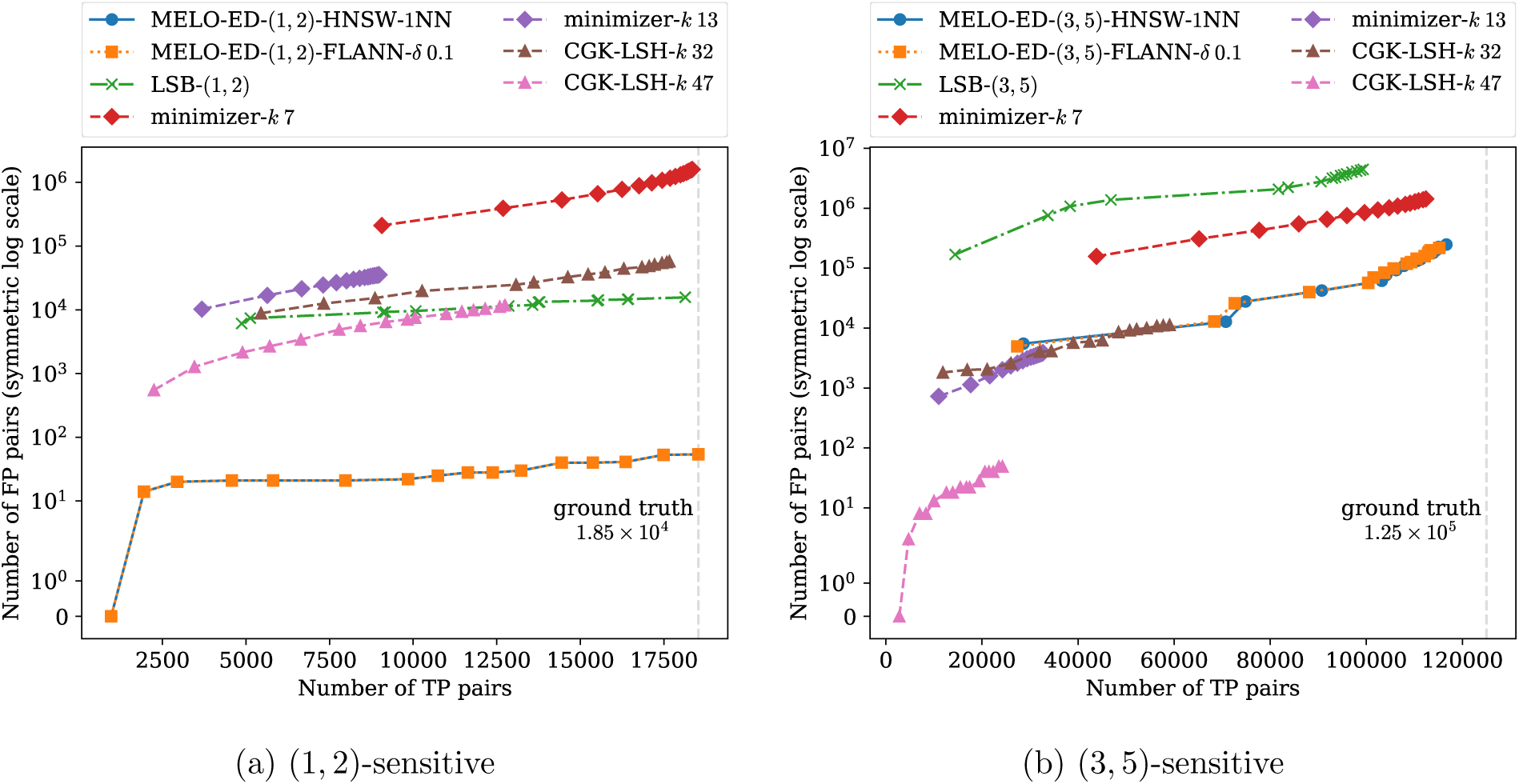
Results of the barcode experiment. In subfigure (a), the HNSW and the FLANN indexes produce identical results which yield overlapping curves.

Consistent with previous experiments, MELO-ED significantly outperforms all competing methods on this real dataset, achieving near-perfect recall while maintaining high precision. This demonstrates that MELO-ED can recover almost all previously discarded reads with minimal false positives, making it a practical and cost-effective addition to single-cell sequencing pipelines.

Figure 10 reports the running times of the evaluated methods on this task. Observe that MELO-ED using the HNSW index achieves running times comparable to traditional seeding- and embedding-based methods, while delivering substantially improved accuracy. Moreover, its running time is insensitive to the similarity thresholds (*d*_1_*, d*_2_), making it a versatile tool suitable for a range of application-specific requirements. The FLANN index exhibits significantly higher running times, primarily due to the internal procedure of fitting an optimal parameter configuration for the input dataset. In practical settings where the base set remains fixed, this overhead represents a one-time cost that can be amortized over multiple query batches.

**Figure 10:**
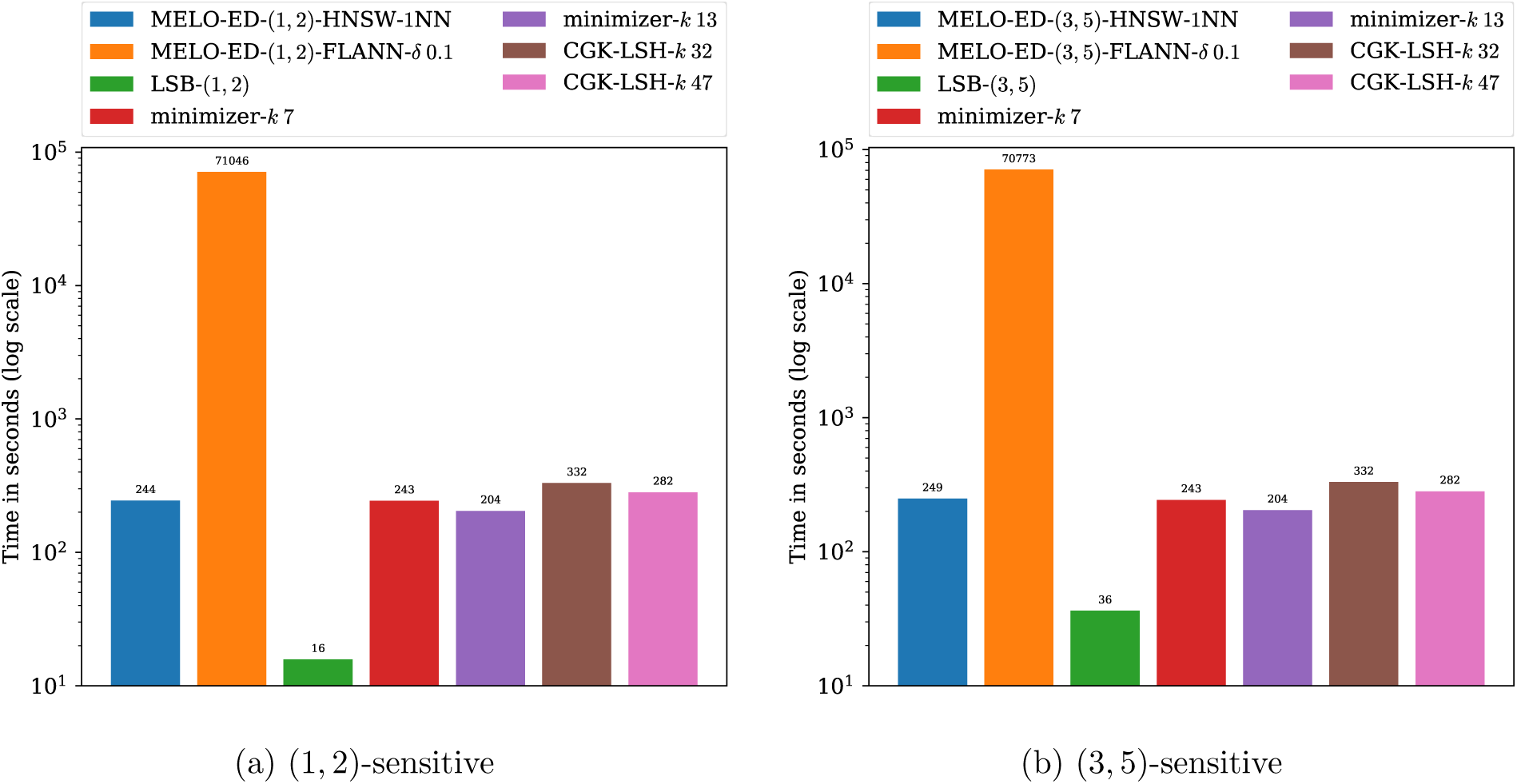
Comparison of running times for the barcode classification experiment.

## 5 Conclusion and Discussion

We present MELO-ED, a new embedding scheme for the edit distance. Unlike existing embedding methods that generate a single vector, which is known to suffer from a large distortion for the edit distance, MELO-ED overcomes this fundamental limitation by allowing for multiple embeddings. Unlike the learned LSB method, which maps each sequence to buckets of hash codes, MELO-ED generates continuous embeddings, providing stronger guarantees that one or a few of them can capture fine-grained similarities between sequences.

Methodologically, inception-style CNN layers enable MELO-ED to simultaneously detect multi-scale patterns, learning both local edit operations and the global contextual organization of nucleotide sequences. Using a Siamese framework trained with a hinge contrastive loss, the network explicitly optimizes for locality-sensitive properties under edit distance, ensuring embeddings stay discriminative especially at category boundaries where edits change classification status.

Experimental results demonstrated that MELO-ED achieved greater separation capability across various (*d*_1_,*d*_2_) thresholds, especially for long sequences and difficult-to-distinguish threshold ranges, such as *N* = 20, *d*_1_*, d*_2_ = (3, 4) and *N* = 100, *d*_1_*, d*_2_ = (5, 6). In many settings, MELO-ED achieves near-perfect accuracy, demonstrating its expressive power. Combined with an indexing approach, MELO-ED substantially outperforms competing methods in sequence neighbor search and in barcode classification. MELO-ED presents a major step forward in large-scale sequence comparison for edit distance and we expect its high impact in the field.

One limitation of MELO-ED is that it requires the two sequences to have the same length. Developing more general or adaptive LSME functions capable of processing variable-length inputs is an important direction of our future work.

## 6 Acknowledgments

This work is supported by the US National Science Foundation (2145171 to M.S.) and by the US National Institutes of Health (R01HG011065 to M.S.).

## References

[1] Arturs Backurs and Piotr Indyk. Edit distance cannot be computed in strongly subquadratic time (unless seth is false). In Proceedings of the forty-seventh annual ACM symposium on Theory of computing, pages 51–58, 2015.

[2] Konstantin Berlin, Sergey Koren, Chen-Shan Chin, James P Drake, Jane M Landolin, and Adam M Phillippy. Assembling large genomes with single-molecule sequencing and locality-sensitive hashing. Nature Biotechnology, 33(6):623–630, 2015.

[3] Diptarka Chakraborty, Elazar Goldenberg, and Michal Kouckỳ. Streaming algorithms for embedding and computing edit distance in the low distance regime. In Proceedings of the 48th ACM Symposium on Theory of Computing (STOC’16), pages 712–725, 2016.

[4] Zhihao Chang, Linzhu Yu, Yanchao Xu, and Wentao Hu. Neural embeddings for knn search in biological sequence. In Proceedings of the AAAI Conference on Artificial Intelligence, volume 38, No. 1, pages 38–45, 2024.

[5] Ke Chen and Mingfu Shao. Locality-sensitive bucketing functions for the edit distance. Algorithms for Molecular Biology, 18(1):7, 2023.

[6] Gabriele Corso, Zhitao Ying, Michal Pándy, Petar Veličković, Jure Leskovec, and Pietro Liò. Neural distance embeddings for biological sequences. Advances in Neural Information Processing Systems, 34:18539–18551, 2021.

[7] Xinyan Dai, Xiao Yan, Kaiwen Zhou, Yuxuan Wang, Han Yang, and James Cheng. Convolutional embedding for edit distance. In proceedings of the 43rd international ACM SIGIR conference on Research and Development in information retrieval, pages 599–608, 2020.

[8] Mayur Datar, Nicole Immorlica, Piotr Indyk, and Vahab S Mirrokni. Locality-sensitive hashing scheme based on p-stable distributions. In Proceedings of the twentieth annual symposium on Computational geometry, pages 253–262, 2004.

[9] Thomas Dencker, Chris-André Leimeister, Michael Gerth, Christoph Bleidorn, Sagi Snir, and Burkhard Morgenstern. ‘multi-spam’: a maximum-likelihood approach to phylogeny reconstruction using multiple spaced-word matches and quartet trees. NAR Genomics and Bioinformatics, 2(1):lqz013, 2020.

[10] Matthijs Douze, Alexandr Guzhva, Chengqi Deng, Jeff Johnson, Gergely Szilvasy, Pierre-Emmanuel Mazaré, Maria Lomeli, Lucas Hosseini, and Hervé Jégou. The faiss library. IEEE Transactions on Big Data, 2025.

[11] Nan Du, Jiao Chen, and Yanni Sun. Improving the sensitivity of long read overlap detection using grouped short *k*-mer matches. BMC Genomics, 20(2):49–62, 2019.

[12] Robert Edgar. Syncmers are more sensitive than minimizers for selecting conserved k-mers in biological sequences. PeerJ, 9:e10805, 2021.

[13] Mahdie Eghdami, Mahmoud Naghibzadeh, and Hamid Noori. Accelerating long-read overlap detection for genome assembly with a two-hash mtable strategy. Computational Biology and Chemistry, page 108576, 2025.

[14] Huan Fan, Anthony R Ives, Yann Surget-Groba, and Charles H Cannon. An assembly and alignment-free method of phylogeny reconstruction from next-generation sequencing data. BMC genomics, 16(1):522, 2015.

[15] Giulia Guidi, Marquita Ellis, Daniel Rokhsar, Katherine Yelick, and Aydın Buluç. Bella: Berkeley efficient long-read to long-read aligner and overlapper. In SIAM Conference on Applied and Computational Discrete Algorithms (ACDA21), pages 123–134. SIAM, 2021.

[16] Piotr Indyk and Rajeev Motwani. Approximate nearest neighbors: towards removing the curse of dimensionality. In Proceedings of the thirtieth annual ACM symposium on Theory of computing, pages 604–613, 1998.

[17] Amir Joudaki, Alexandru Meterez, Harun Mustafa, Ragnar Groot Koerkamp, André Kahles, and Gunnar Rätsch. Aligning distant sequences to graphs using long seed sketches. Genome Research, 33(7):1208–1217, 2023.

[18] Amir Joudaki, Gunnar Ratsch, and André Kahles. Fast alignment-free similarity estimation by tensor sketching. bioRxiv, 2020.

[19] Robert Krauthgamer and Yuval Rabani. Improved lower bounds for embeddings into *l*_1_. SIAM Journal on Computing, 38(6):2487–2498, 2009.

[20] Heng Li. Minimap2: pairwise alignment for nucleotide sequences. Bioinformatics, 34(18):3094–3100, 2018.

[21] Xiang Li, Ke Chen, and Mingfu Shao. Efficient seeding for error-prone sequences with subse-qhash2. Bioinformatics, 41(8):btaf418, 2025.

[22] Xiang Li, Qian Shi, Ke Chen, and Mingfu Shao. Seeding with minimized subsequence. Bioinformatics, 39(Supplement 1):i232–i241, 06 2023.

[23] Junwei Luo, Ranran Chen, Xiaohong Zhang, Yan Wang, Huimin Luo, Chaokun Yan, and Zhanqiang Huo. Lrod: an overlap detection algorithm for long reads based on k-mer distribution. Frontiers in Genetics, 11:632, 2020.

[24] Benjamin Dominik Maier and Kristoffer Sahlin. Entropy predicts fuzzy-seed sensitivity. bioRxiv, page 2022.10.13.512198, 2022.

[25] Guillaume Maŗcais, Dan DeBlasio, and Carl Kingsford. Asymptotically optimal minimizers schemes. Bioinformatics, 34(13):i13–i22, 2018.

[26] Guillaume Maŗcais, Dan DeBlasio, Prashant Pandey, and Carl Kingsford. Locality-sensitive hashing for the edit distance. Bioinformatics, 35(14):i127–i135, 2019.

[27] Marius Muja and David G Lowe. Fast approximate nearest neighbors with automatic algorithm configuration. VISAPP (1), 2(331-340):2, 2009.

[28] Marius Muja and David G Lowe. Fast matching of binary features. In 2012 Ninth conference on computer and robot vision, pages 404–410. IEEE, 2012.

[29] Marius Muja and David G Lowe. Scalable nearest neighbor algorithms for high dimensional data. IEEE transactions on pattern analysis and machine intelligence, 36(11):2227–2240, 2014.

[30] Brian D Ondov, Todd J Treangen, Páll Melsted, Adam B Mallonee, Nicholas H Bergman, Sergey Koren, and Adam M Phillippy. Mash: fast genome and metagenome distance estimation using minhash. Genome biology, 17(1):132, 2016.

[31] Rafail Ostrovsky and Yuval Rabani. Low distortion embeddings for edit distance. Journal of the ACM (JACM), 54(5):23–es, 2007.

[32] Michael Roberts, Wayne Hayes, Brian R Hunt, Stephen M Mount, and James A Yorke. Reducing storage requirements for biological sequence comparison. Bioinformatics, 20(18):3363–3369, 2004.

[33] Michael Roberts, Brian R Hunt, James A Yorke, Randall A Bolanos, and Arthur L Delcher. A preprocessor for shotgun assembly of large genomes. Journal of Computational Biology, 11(4):734–752, 2004.

[34] Kristoffer Sahlin. Effective sequence similarity detection with strobemers. Genome Research, 31(11):2080–2094, 2021.

[35] Kristoffer Sahlin. Strobealign: flexible seed size enables ultra-fast and accurate read alignment. Genome Biology, 23(1):1–27, 2022.

[36] Shahab Sarmashghi, Kristine Bohmann, M Thomas P. Gilbert, Vineet Bafna, and Siavash Mirarab. Skmer: assembly-free and alignment-free sample identification using genome skims. Genome biology, 20(1):34, 2019.

[37] Saul Schleimer, Daniel S Wilkerson, and Alex Aiken. Winnowing: local algorithms for document fingerprinting. In Proceedings of the 2003 ACM SIGMOD International Conference on Management of Data (SIGMOD/PODS’03), pages 76–85, 2003.

[38] Gregory E. Sims, Se-Ran Jun, Guohong A. Wu, and Sung-Hou Kim. Alignment-free genome comparison with feature frequency profiles (ffp) and optimal resolutions. Proceedings of the National Academy of Sciences, 106(8):2677–2682, 2009.

[39] Yan Song, Haixu Tang, Haoyu Zhang, and Qin Zhang. Overlap detection on long, error-prone sequencing reads via smooth q-gram. Bioinformatics, 36(19):4838–4845, 2020.

[40] Fang Wang, Yibin Wang, Xiaofei Zeng, Shengcheng Zhang, Jiaxin Yu, Dongxi Li, and Xingtan Zhang. Mike: an ultrafast, assembly-, and alignment-free approach for phylogenetic tree construction. Bioinformatics, 40(4):btae154, 2024.

[41] Anuradha Wickramarachchi and Yu Lin. Metagenomics binning of long reads using read-overlap graphs. In RECOMB International Workshop on Comparative Genomics, pages 260–278. Springer, 2022.

[42] Yupei You, YD Prawer, Ricardo De Paoli-Iseppi, C. P Hunt, C. L. Parish, H Shim, and M. B. Clark. Identification of cell barcodes from long-read single-cell rna-seq with blaze. Genome Biology, 24(1):66, 2023.

[43] Xin Yuan, Ke Chen, Xiang Li, Qian Shi, and Mingfu Shao. Learning locality-sensitive bucketing functions. Bioinformatics, 40(Supplement 1):i318–i327, 2024.

[44] Xiyuan Zhang, Yang Yuan, and Piotr Indyk. Neural embeddings for nearest neighbor search under edit distance, 2020.

