## Supplemental Figures for "MELO-ED: learning locality-sensitive multi-embeddings for edit distance"

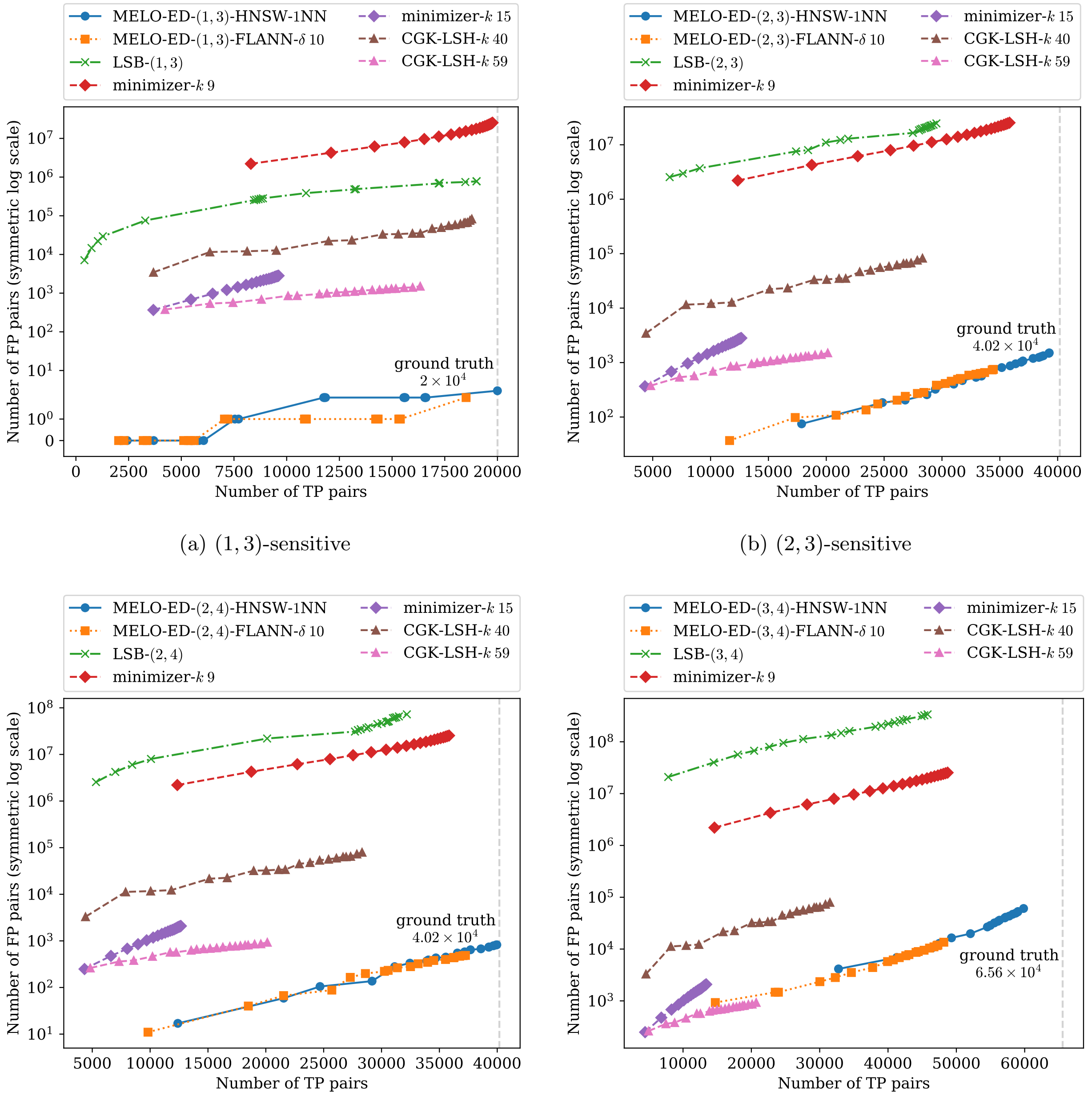

(a) $(1, 3)$-sensitive

(b) $(2, 3)$-sensitive

(c) $(2, 4)$-sensitive

(d) $(3, 4)$-sensitive

**Supplementary Figure 1**: Additional results of applying a neighbor search index to the embeddings produced by MELO-ED on the test dataset, compared with other methods.

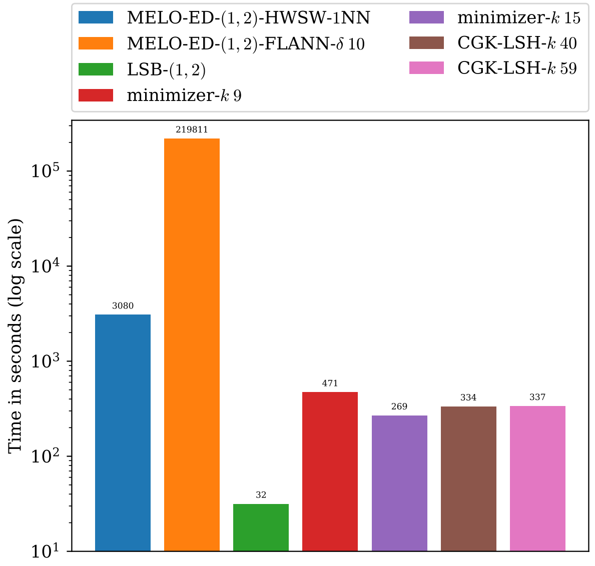

(a) $(1,2)$-sensitive

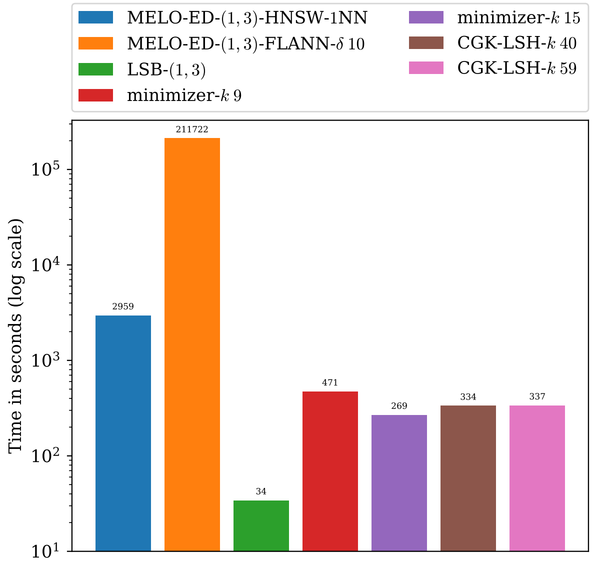

(b) $(1,3)$-sensitive

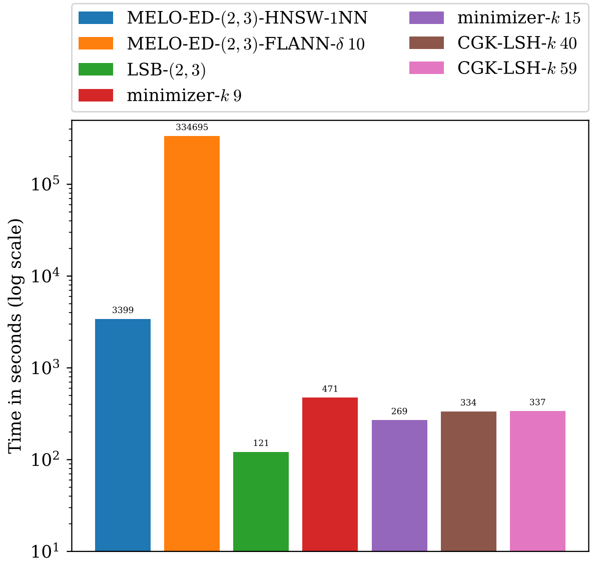

(c) $(2,3)$-sensitive

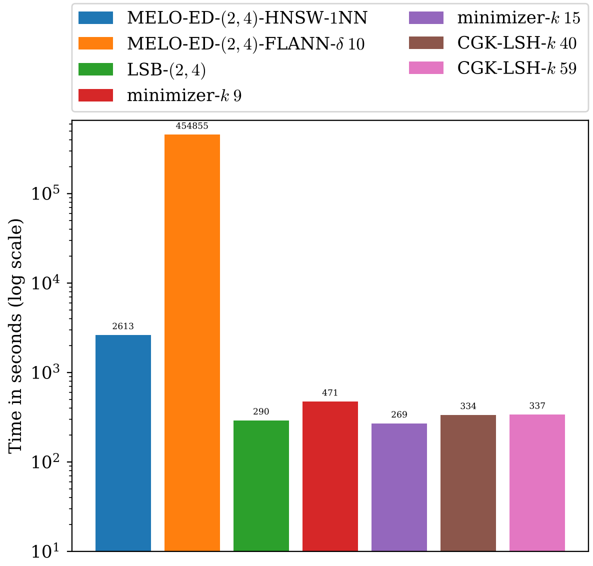

(d) $(2,4)$-sensitive

**Supplementary Figure 2**: Time of applying a neighbor search index to the embeddings produced by MELO-ED on the test dataset, compared with other methods.

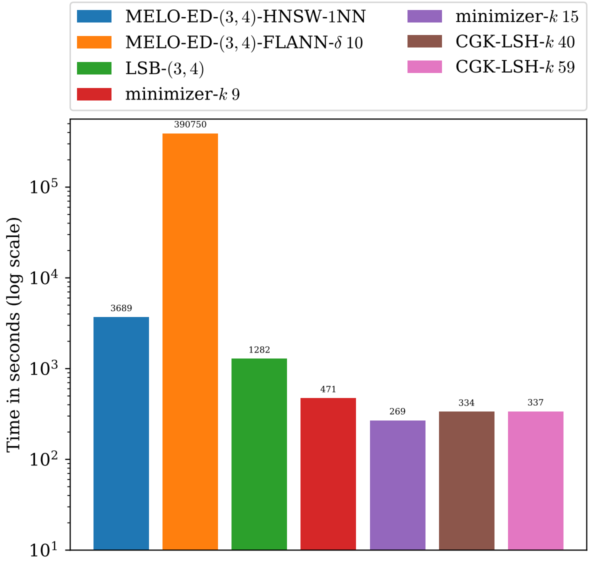

(a) $(3, 4)$-sensitive

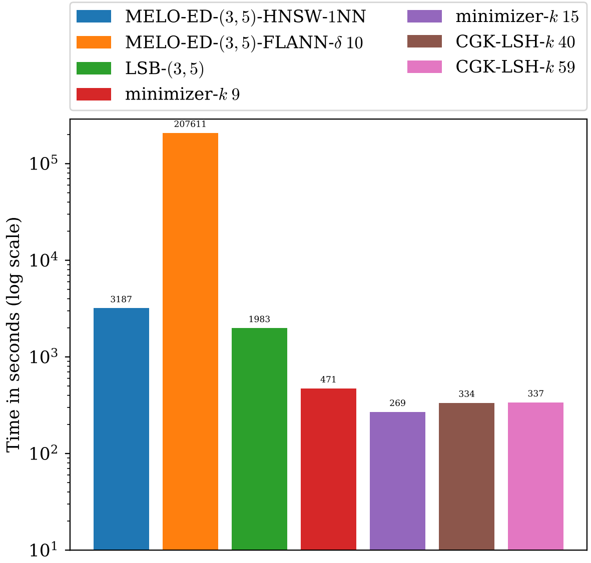

(b) $(3, 5)$-sensitive

**Supplementary Figure 3**: (Cont'd) Time of applying a neighbor search index to the embeddings produced by MELO-ED on the test dataset, compared with other methods.

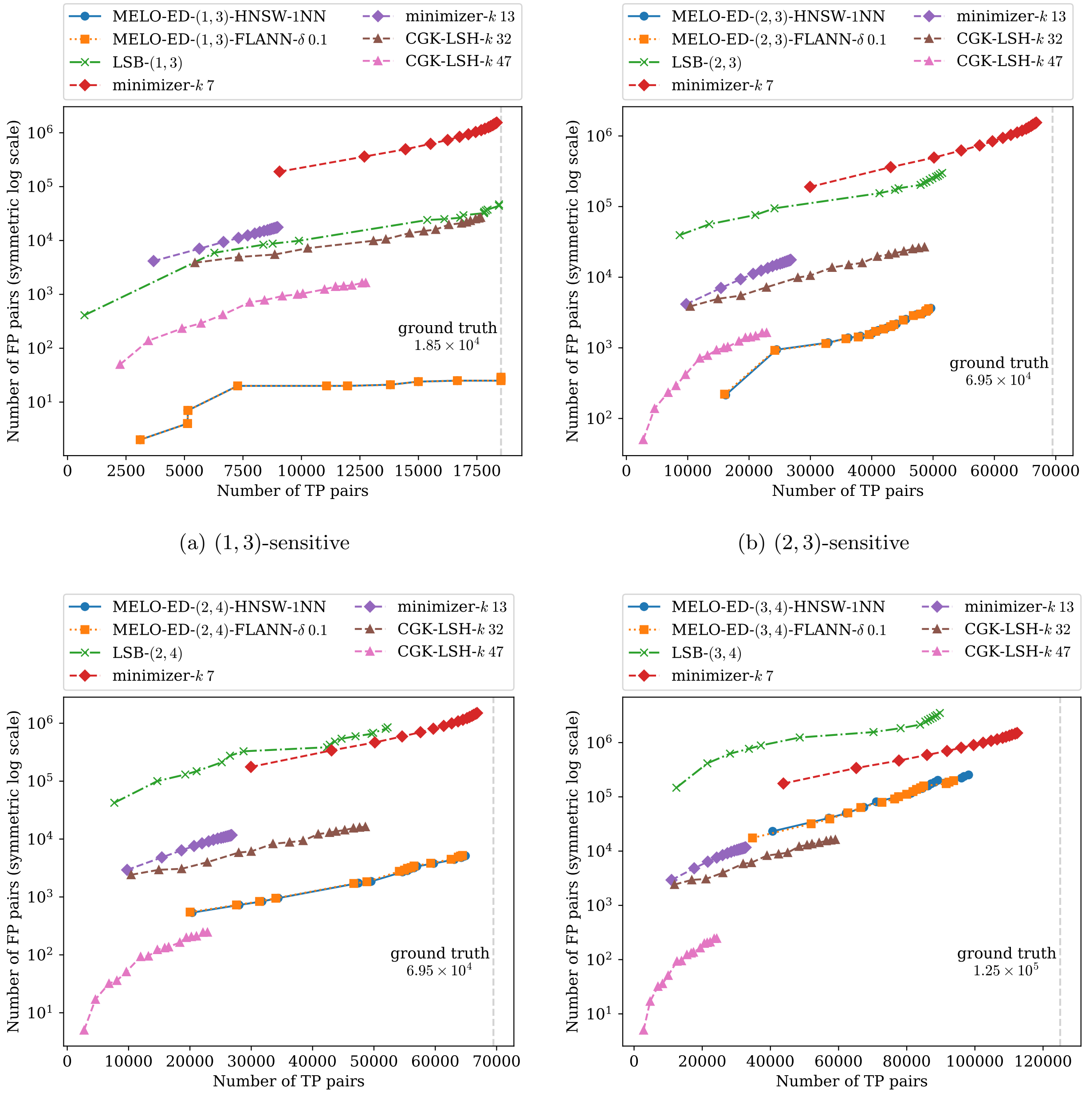

(a) $(1, 3)$-sensitive

(b) $(2, 3)$-sensitive

(c) $(2, 4)$-sensitive

(d) $(3, 4)$-sensitive

**Supplementary Figure 4**: Additional results of applying a neighbor search index to the embeddings produced by MELO-ED on the barcode experiment.

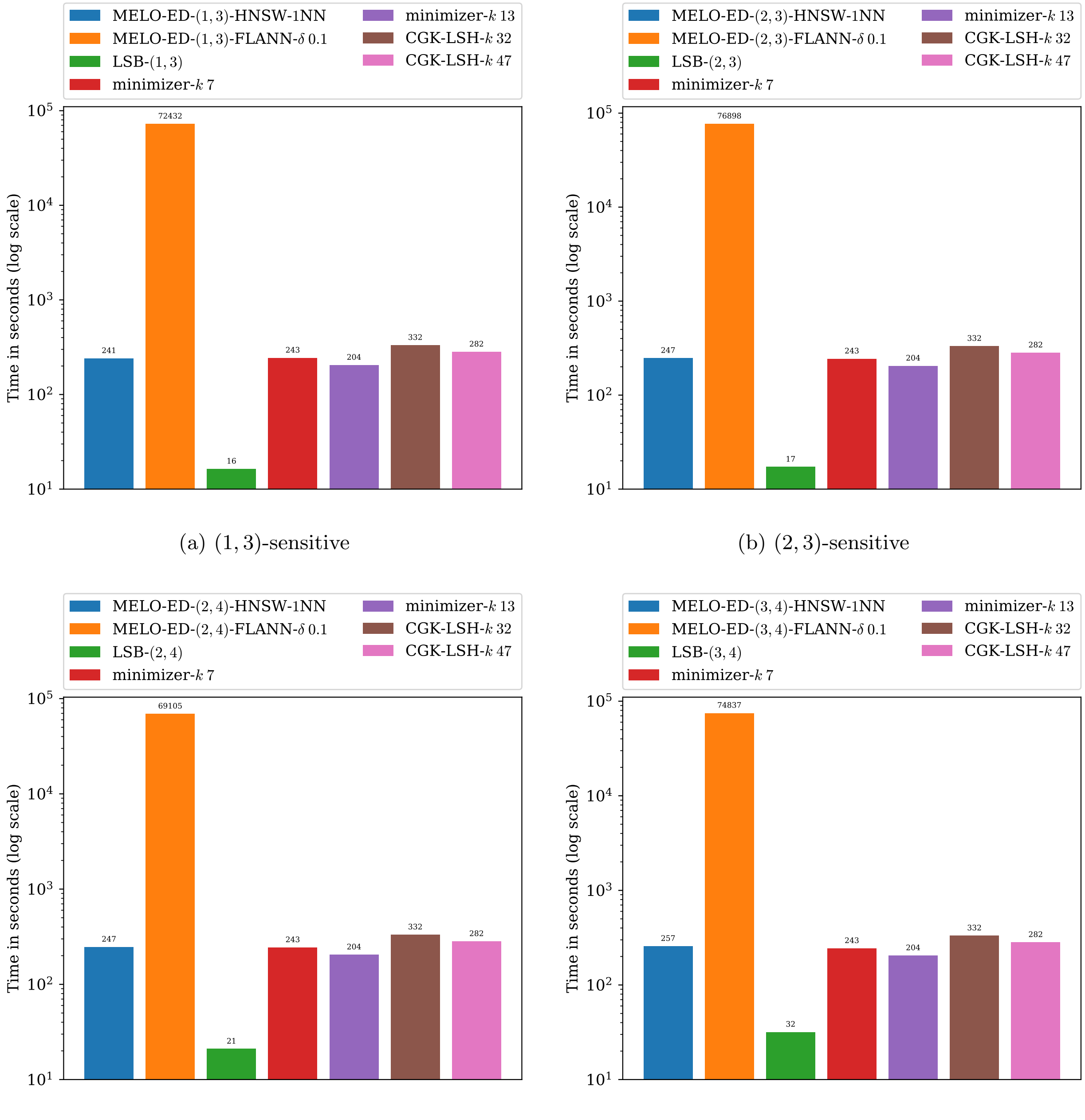

(a) $(1, 3)$-sensitive

(b) $(2, 3)$-sensitive

(c) $(2, 4)$-sensitive

(d) $(3, 4)$-sensitive

**Supplementary Figure 5**: Time of applying a neighbor search index to the embeddings produced by MELO-ED on the barcode experiment.

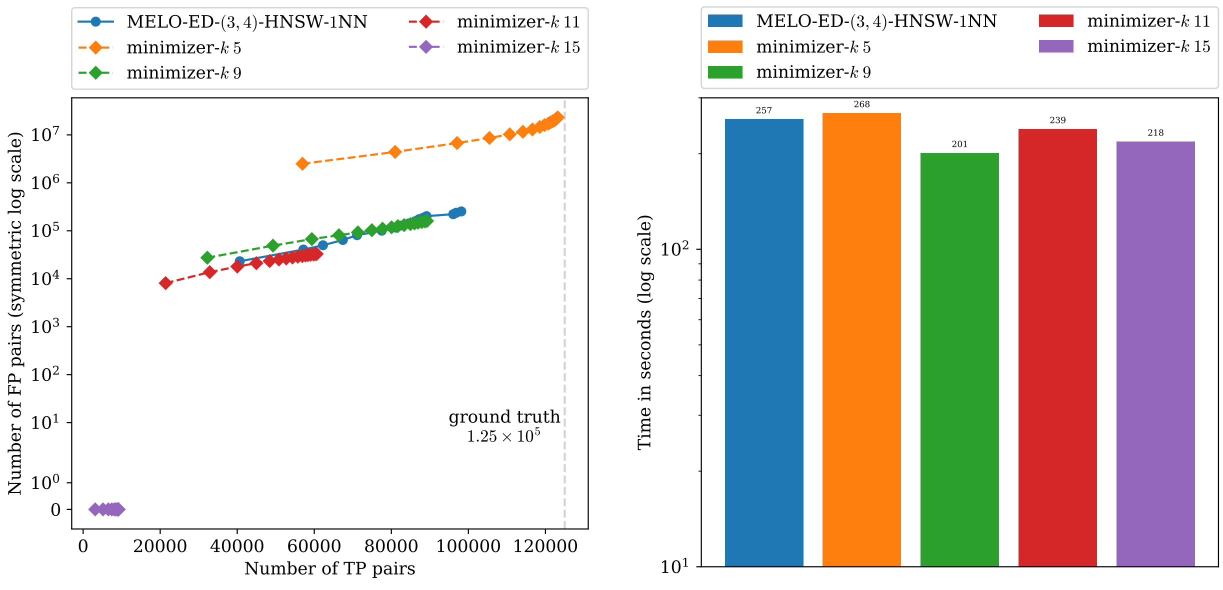

(a) $(3, 4)$-sensitive

(b) $(3, 4)$-sensitive

**Supplementary Figure 6**: Additional results and running times for the barcode experiment. Since MELO-ED outperforms the minimizer methods for all tested $(d_1, d_2)$ values except the $(3, 4)$-sensitive setting, we include additional minimizer results with varied parameter $k$ for the $(3, 4)$-sensitive case. These plots show that MELO-ED still attains higher recall at comparable levels of false positives and running time.
